# Localized cell-surface sampling of a secreted factor using cell-targeting beads

**DOI:** 10.1101/2020.06.11.147140

**Authors:** Tammi L. van Neel, Samuel B. Berry, Erwin Berthier, Ashleigh B. Theberge

**Affiliations:** Department of Chemistry, University of Washington, Box 351700, Seattle, Washington 98195, USA; Department of Urology, University of Washington, Seattle, Washington 98105, USA

## Abstract

Intercellular communication through the secretion of soluble factors plays a vital role in a wide range of biological processes (e.g., homeostasis, immune response), yet identification and quantification of many of these factors can be challenging due to their degradation or sequestration in cell culture media prior to analysis. Here, we present a customizable bead-based system capable of simultaneously binding to live cells (through antibody-mediated cell-tethering) and capturing cell-secreted molecules. Our functionalized beads capture secreted molecules (e.g., hepatocyte growth factor secreted by fibroblasts) that are diminished when sampled *via* traditional supernatant analysis techniques (p < 0.05), effectively rescuing reduced signal in the presence of neutralizing components in the cell culture media. Our system enables capture and analysis of molecules integral to chemical communication that would otherwise be markedly decreased prior to analysis.

**For Table of Contents Only:** 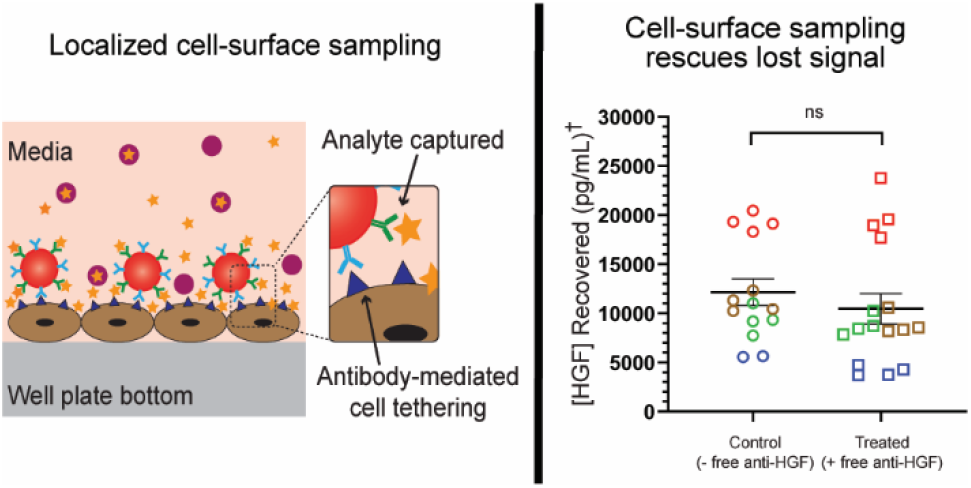

## Introduction

Chemical signaling events involving cell-secreted soluble factors (e.g., growth factors, cytokines) play a vital role in the induction of cellular functions and larger systematic biological processes including immune response, maintenance of homeostasis, and cellular differentiation.^1^ However, detecting these molecules to decipher this complex signaling landscape is often hindered through the degradation, sequestration, or neutralization of important signaling molecules by extracellular factors such as enzymes or receptors.^2,3^ The elimination of these short-lived soluble factors from a cellular microenvironment is an important component of chemical signaling processes, yet their absence leads to an incomplete snapshot of the signaling microenvironment, as these transient factors cannot be easily analyzed. For example, the instability of eicosanoids (e.g., leukotrienes, prostaglandins)^4,5^ or the degradation of cytokines by proteases^6–8^ or poor storage conditions in clinical settings^9^ makes their quantification challenging, hindering understanding of their role in biological processes such as inflammation and cancer. Identification and quantification of these key short-lived factors in the context of their localized signaling milieu can provide important insight into the signaling mechanisms that mediate biological processes within complex *in vivo* and *in vitro* systems.^10,11^

Various analytical and quantification methods, such as mass spectrometry and enzyme-linked immunosorbent assays (ELISA), have been developed that examine cell culture supernatants (i.e., conditioned media) or biological fluids (e.g., serum, urine) to provide important information on the makeup of cellular secretion profiles.^12^ However, these methods often rely on sampling processes wherein important effector molecules may be degraded, sequestered, or converted on time scales faster than those required for sample preparation and analysis, resulting in diminished signal; further, these readouts are typically used as end-point analyses that lack the temporal resolution provided by *in situ* methods and analyses.^13^ More targeted approaches that integrate sample collection and readout, such as compartmentalized microfluidic cell culture platforms for *in situ* bead-based assays^14,15^, integrated microchip single-cell culture and analysis devices^16^, small-volume cell-encapsulation and -sensor systems^17–21^ and enzyme-linked immunosorbent spot (ELISpot) assays^22,23^, address the limitations posed by traditional techniques and enable precise *in situ* analysis of culture systems at flexible timepoints throughout the experiment. These integrated culture and analysis platforms allow users to probe specific phenomena using systems with excellent spatial and temporal detection resolution, as well as single-cell resolution for secretome analysis. However, many of these platforms require complex platforms and advanced fabrication facilities, decreasing their transferability, and rely on materials such as polydimethylsiloxane (PDMS), which has been shown to absorb small molecules.^24,25^ Therefore, we sought to add to the analytical capabilities demonstrated in these advanced technologies through the creation of a transferable and easily-deployed system compatible with virtually all culture setups and sizes.

Bead-based technologies have been widely used for both the analysis of soluble factors within biological samples (e.g., bead-based ELISA) and to selectively capture and analyze cells from a mixed culture (e.g., magnetic bead-based cell isolation).^26–28^ Additionally, the use of beads for targeted cellular secretome analysis has been applied in customized systems (often at a single-cell resolution) in applications including T cell secretion and function in cancer^15,21,29^ and B cell secretion of antibodies for vaccination and immunity.^19,20^ However, for larger scale applications (i.e., greater than single cell or microfluidic analyses), there lacks a broadly deployable bead-based technology to examine the production of transient soluble factors and the origin of those factors (Figure 1); additionally, to our knowledge, the ability to simultaneously capture these transient soluble factors and the cell itself with the same bead has not been demonstrated. Here, we introduce a customized, off-the-shelf bead-based approach to enable capture of short lived or unavailable compounds from within existing cell culture systems that can then be coupled with downstream analytical methods such as immunoassays (Figure 1). Our platform consists of a dual-functionalized (DF) magnetic bead with two distinct antibodies, enabling simultaneous cell-binding and signal capture (Figure 1 Bii). Through cell tethering, our DF beads can target a specific cell type *via* cell-specific surface markers, as well as capture cell-secreted signals before they enter the bulk solution, where they may be sequestered or degraded. Here, we demonstrate that our DF beads capture a cell-secreted signal (hepatocyte growth factor, HGF) localized near the cell surface from live fibroblast cultures in the presence of a neutralization factor; in contrast HGF levels are markedly diminished when collected through traditional supernatant analysis. We envision these dual-functionalized beads being employed in a wide range of mono- and multi-cultures, enabling researchers to easily “listen” to cellular communication between different cell populations *in situ* without needing to modify their culture protocols or setup.

**Figure 1.**
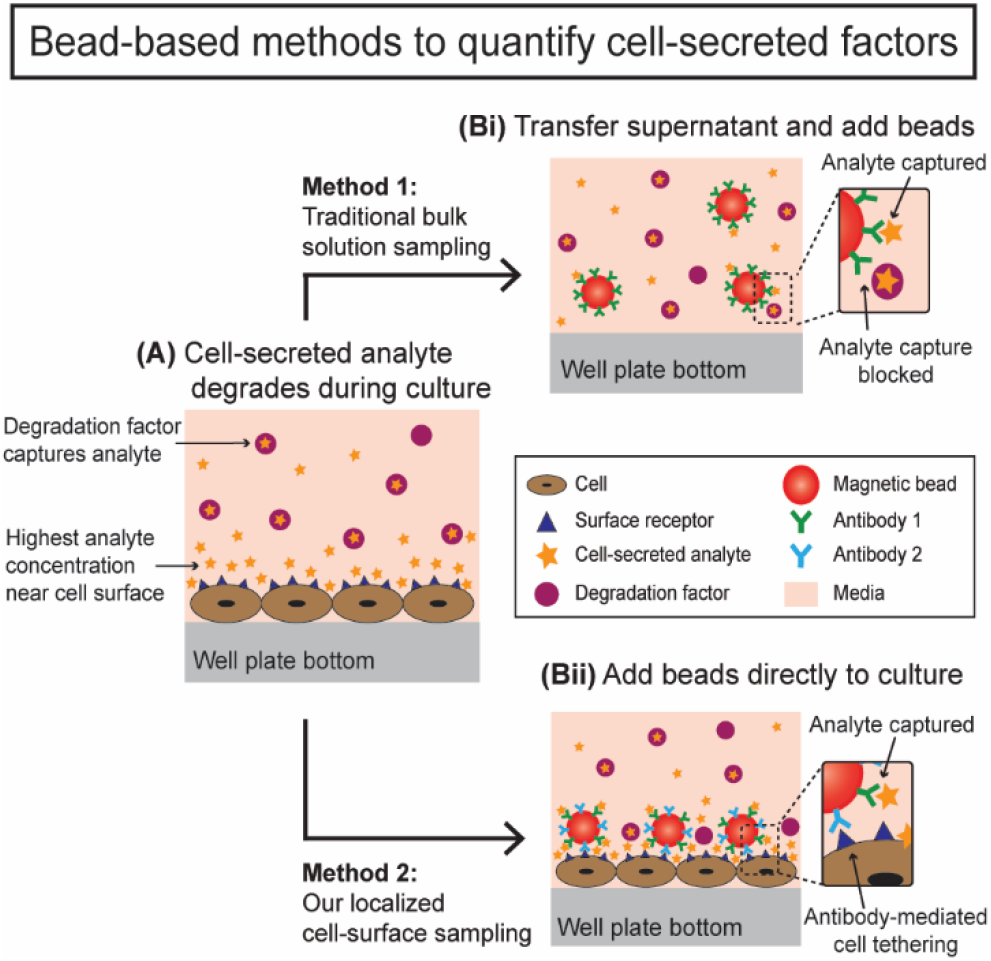
Bead-based approaches to quantifying cell-secreted factors. (A) Cells secrete soluble factors for communication, with the highest concentration immediately after secretion being near the cell surface. (Bi) Traditional methods transfer the cell culture supernatant to another well plate and use beads for bulk solution analyte sampling while (Bii) our new approach utilizes cell-targeting beads in situ for analyte sampling.

## Results and Discussion

To probe the myriad of diverse environments where intercellular signaling occurs, an adaptable platform capable of being deployed across a wide range of signaling landscapes is required. Herein, we present a dual-functionalized (DF) bead system that binds to cells within a live *in vitro* cell culture system to analyze a secreted factor of interest in the presence and absence of a factor that may degrade or neutralize, or sequester the factor of interest. As a proof-of-concept model, we selected neonatal normal human dermal fibroblasts (NHDFn) expressing surface marker CD90 (Figure S1) and secreting hepatocyte growth factor (HGF) as our cellular and molecular target, respectively. HGF is a vital factor in multiple biological processes, including organ development, liver repair, and cancer progression, underscoring the importance of fibroblast-driven chemical communication in a range of biological settings.^30^ We designed and created dual-functionalized MagPlex beads covalently bound with antibodies against CD90 (anti-CD90) and HGF (anti-HGF) to target the cell surface marker and secreted factor, respectively (Figure S2).^31^ To validate that the DF beads can successfully capture HGF at biologically relevant concentrations while simultaneously binding to the cell surface, we conducted two independent cell-binding and analyte capture assays (Figure 3-4, S3-7).

First, we performed a standard sandwich immunoassay using our DF beads (anti-CD90 and anti-HGF) and commercially available mono-functionalized (MF, anti-HGF only) beads with spiked HGF to compare HGF detection capabilities using the DF beads and MF beads when used according to traditional protocols (i.e., supernatant analysis). Both DF and MF beads successfully captured spiked HGF at a range of concentrations, enabling fitting to a five-parameter logistic (5PL) calibration curve as recommended by the manufacturer^31^ and supporting the use of the DF beads as a tool for capturing and measuring secreted factors from cells (Figure 2). Differences in the curve shape is due to variation of kit performance; the kit performance variation was within accepted variability ranges based on the manufacturer’s standards and protocols. We observed different limits of detection (LODs) between the two beads consistent with the different bead functionalization: DF beads, 104.25 pg/mL; MF beads, 34.87 pg/mL (Figure 2). We also observed that DF beads had a 17-36% lower total fluorescent signal than the MF beads (Figure 2). This is expected, as the number of available binding sites on the surface of the DF beads is less than the binding sites on the surface of the MF beads due to the presence of anti-CD90 antibodies, thereby decreasing the total number of fluorescent detection antibodies that will bind to HGF on the surface of the DF bead.

**Figure 2:**
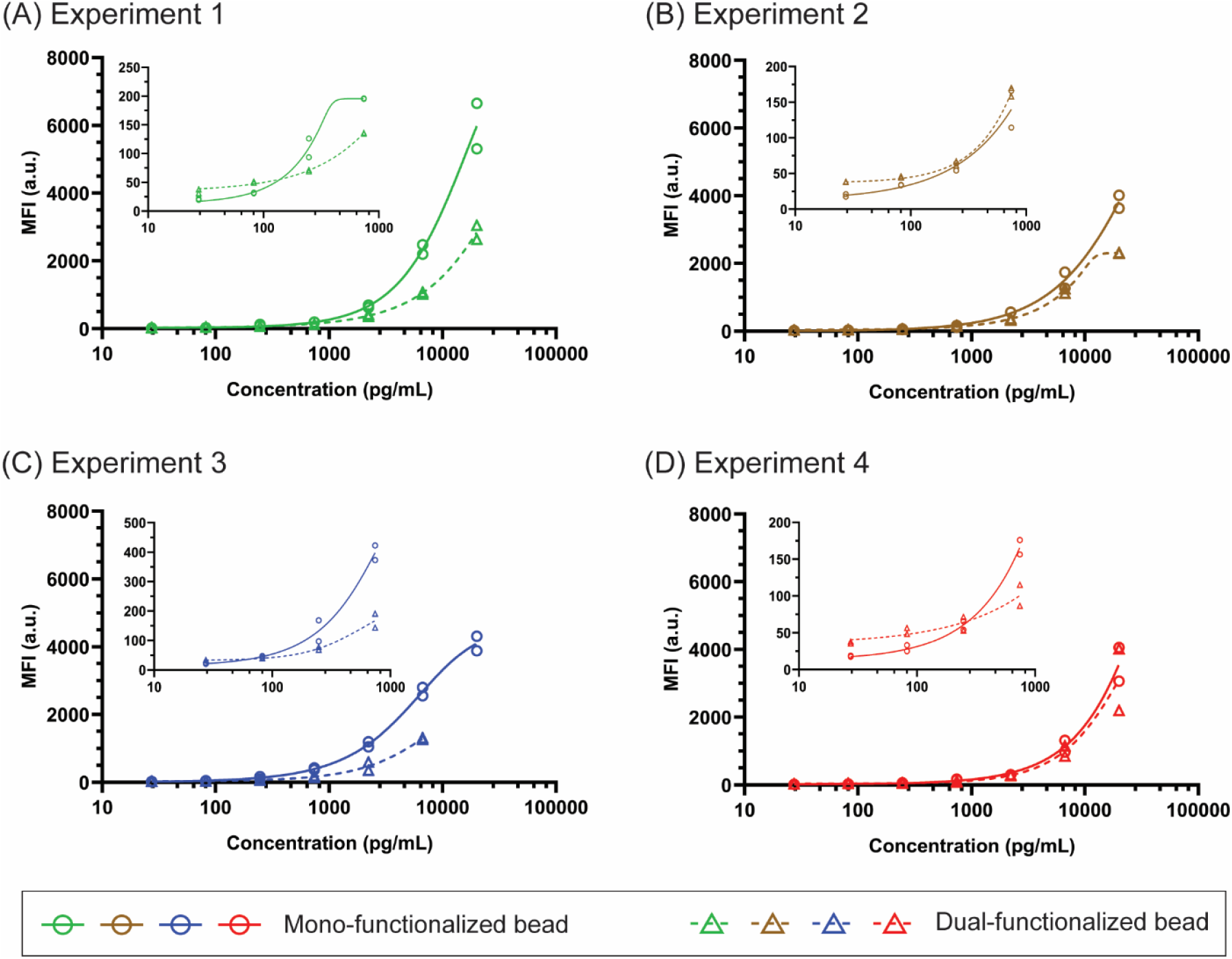
Calibration curves for mono-(MF) and dual-functionalized (DF) beads demonstrate ability of both beads to capture HGF. Per MagPlex manufacturer’s recommendation, a five-parameter logistic fit was used to fit calibration curves and demonstrate the HGF-bead binding ability for each independent experiment (A: Experiment 1; B: Experiment 2; C: Experiment 3; D: Experiment 4) in duplicate (duplicates are shown as two plotted circles or triangles). Curves follow the expected trend based on manufacturer’s technical notes.^31^ Decreased signal is observed for DF beads as the surface has roughly half the binding sites for HGF compared to MF beads. Insets are of first four points of the calibration curves. All curves are plotted on a logarithmic scale for visualization.

Second, we performed a cell-binding assay wherein we incubated DF beads with CD90-expressing NHDFn cells (Figure 3) and quantified their ability to remain bound to the cell surface after multiple wash steps (Figure S3-S6 and Table S1). Beads were added directly to a 96-well plate culture of NHDFn cells and incubated for 2 hours; after incubation, the culture was rinsed three times to remove unbound or nonspecifically-bound beads. The remaining beads were then imaged and counted to quantify the ability of the DF beads to bind CD90^**+**^cells (i.e., NHDFn) (Figure 3, S3-S7, and Table S1). In addition to the fluorescence microscopy images shown in Figures 3 and S3, phase contrast images at 10X and 20X are provided in Figures S4 and S5. After washing, 71% of the beads were retained, suggesting successful CD90-mediated cell binding (Figure 3, S3, and Table S1). Further optimization of the bead functionalities (i.e., analyte capture and cell binding) can be accomplished by adjusting the ratio of antibodies on the surface, as well as through modulation of the specific biological system of interest to increase expression of cell-binding markers or secretion of soluble factors.

**Figure 3.**
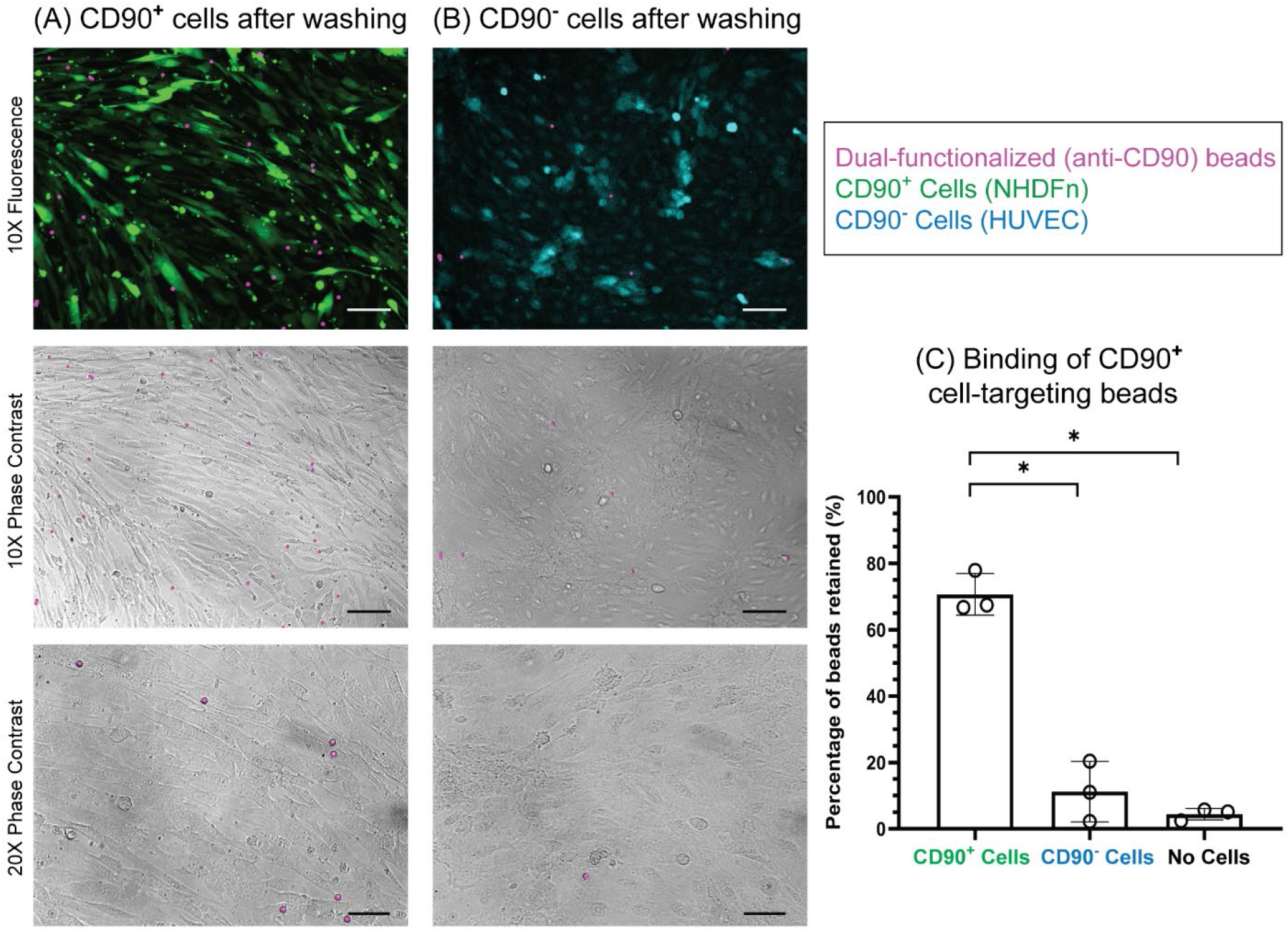
Dual-functionalized (DF) beads (functionalized with anti-CD90 and anti-HGF) target CD90^**+**^cells (NHDFn). (A and B) Representative images of CD90^**+**^cells stained with CellTracker Green (green) and CD90^**−**^cells stained with CellTracker Blue (blue); cells were seeded at a density of 2.6×10^4^ cells/mL, cultured for 3 days, and incubated with DF beads (pink) for 2 h. Images of wells were taken after washing to determine successful bead-binding to the cell surface receptor CD90. (A) CD90^**+**^cells show an increased bead retention compared to (B) CD90^**−**^cells. (C) An average of 71% of DF beads were retained when incubated with CD90^**+**^cells, significantly higher bead retention than for CD90^-^ cells or no cells (empty well). Some nonspecific binding was observed for beads placed in an empty well (<5% of beads retained). Bar graphs represent mean ± SD of n=3 independent experiments. Brown-Forsynthe and Welch ANOVA tests were used followed by Dunnett’s T3 test. * p < 0.05. 10X Scale bar = 100μm. 20X Scale bar = 50μm.

To demonstrate one utility of our DF bead system, we created a model NHDFn culture system where we test the ability of the beads to capture HGF signal before the signal is diminished in the bulk solution through neutralization with free anti-HGF antibodies; we hypothesized that this resuscitation of HGF signal would be achieved through binding of the DF beads to the cell surface so that the beads can capture HGF as it is being secreted from the cell. To test this hypothesis, we cultured NHDFn cells with our DF beads and free anti-HGF antibodies so that secreted HGF could bind to either the beads (and be detected) or the free anti-HGF (and be neutralized); after a 2-hour incubation, the supernatant was collected and the culture was washed to remove any unbound beads or remaining free anti-HGF (Figure 4). The cells were then lysed to release the beads, which were subsequently washed and analyzed using standard Luminex assay protocols.^31^ In parallel, we analyzed the HGF signal captured by MF beads according to the manufacturer’s protocol (i.e., traditional cell culture supernatant sampling) (Figure 4). Following traditional cell culture supernatant sampling protocols, we observed a significant decrease (p < 0.05) in HGF quantification as a result of the presence of free anti-HGF in the cell culture media in comparison to controls without free anti-HGF (Figure 4 Bi), suggesting that a sequestering factor (such as free anti-HGF) is able to reduce the secreted HGF available for quantification in bulk media. In contrast, under identical culture conditions, our dual-functionalized beads deployed on the cell surface restored HGF signal to levels comparable to those found in the absence of free anti-HGF (Figure 4 Bii), suggesting that the DF beads are able to capture signal prior to its reduction in the bulk supernatant. The increased values observed in Experiment 4 are due to biological cell variability which is expected in cell culture experiments. When examining the relative standard deviation, we observed an increased variability in the traditional supernatant sampling (8-62%) while our localized cell-surface sampling method was consistently below 15% (Table S2).

**Figure 4.**
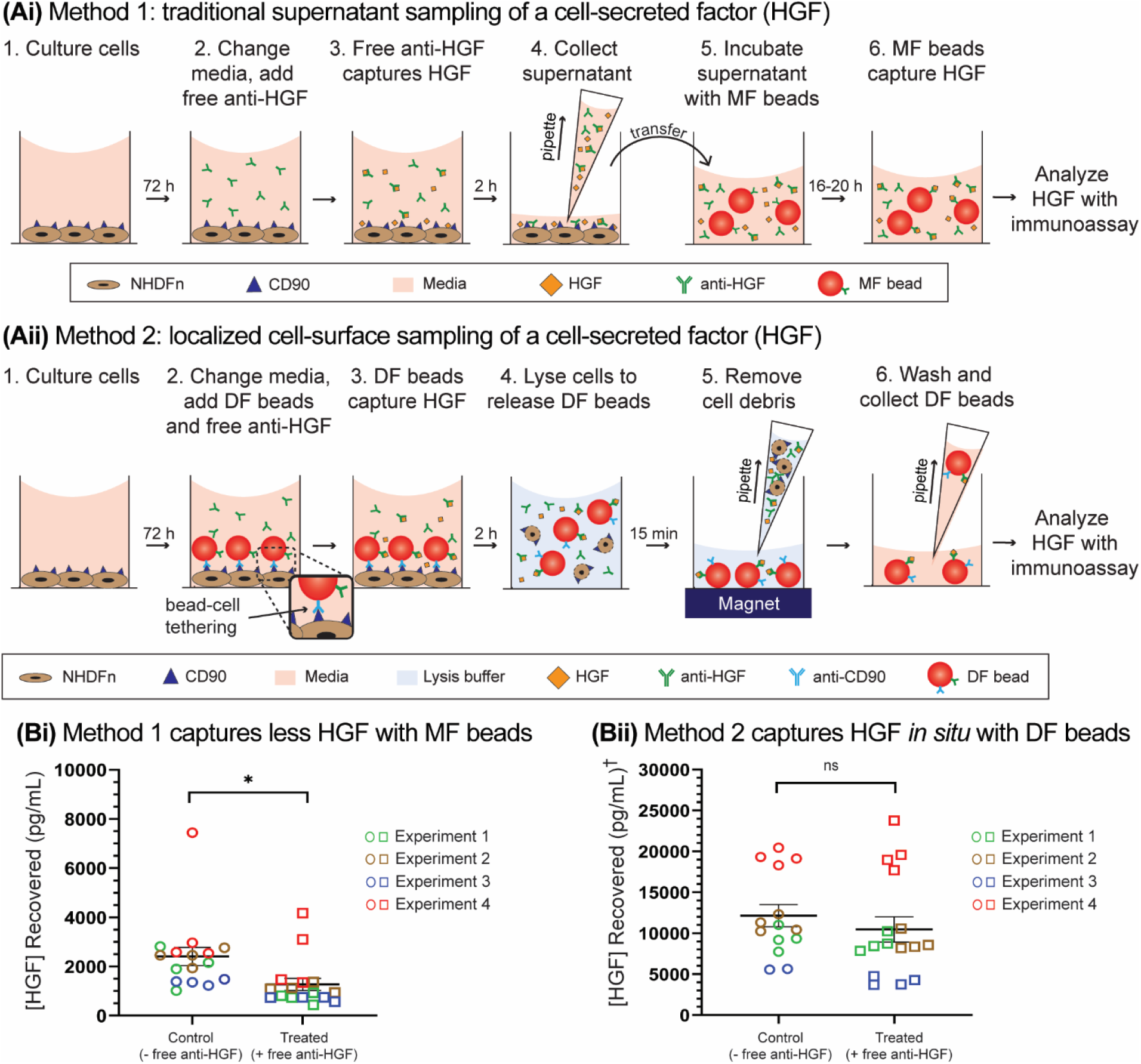
Localized cell-surface sampling recovers HGF signal in the presence of a neutralizing factor. Schematics showing workflow of (Ai) traditional supernatant sampling and (Aii) localized cell-surface sampling of a cell-secreted factor. Mono-functionalized (MF) beads (functionalized with anti-HGF) were used for supernatant sampling while dual-functionalized (DF) beads (functionalized with anti-HGF and anti-CD90) were used for localized cell-surface sampling. (Ai) MF beads are added to supernatant samples collected from cell cultures after 2 h incubation. (Aii) DF beads are added to directly to cultures for a 2 h sampling period. (Bi) Supernatant analysis shows a decrease in recovered HGF when cells are incubated with free anti-HGF whereas (Bii) localized cell-surface sampling with DF beads shows no change in recovered HGF since beads capture HGF before neutralization by free anti-HGF. Error bars are SEM. Experiments were performed in the absence (-) or presence (+) of free anti-HGF and are represented as circles or squares, respectively; each point represents an averaged technical replicate (supernatant sampling) or culture replicate (localized cell-surface sampling). († HGF recovered using DF beads is captured at the surface of the cell, rather than the bulk, leading to increased overall HGF capture. The absolute values in (Bi) should not be directly compared to the absolute values in (Bii), but rather the trend within each graph between control and treated conditions.) Unpaired Student’s t-test. * p = 0.016; ns p = 0.428.

As a control, to verify that the proximity of the DF beads to the cell promotes improved HGF capture, and to eliminate any artifacts associated with cell-bead-binding and the lysis step employed in our localized cell-surface sampling method, we cultured NHDFn cells in close proximity to (10 μm) and at a distance (1.3 mm) from the beads in the presence and absence of free anti-HGF using a Transwell insert setup (Figure S8 A and B). The use of the Transwell insert ensured that the DF beads remained segregated from the cells due to the difference in the membrane pore size and bead diameter (0.2 μm and 5.6 μm, respectively). We found that the beads captured increased levels of HGF (both in the presence and absence of free anti-HGF) when in close proximity to the cells when compared to at a distance. This finding supports that the increased signal capture capabilities of our localized cell-surface sampling method are due to the proximal location of the DF beads on the cell surface (Figure 4, S8 C). While it is likely that MF beads would provide a similar signal if allowed to settle nonspecifically on the cell surface, the ability of the DF beads to specifically bind to cell types of interest enables beads to target the desired cell and ensure the cellular origin of the secreted signal (Figure S7). Further, to validate that the observed HGF signal in our localized cell-surface sampling method (Figure 4 Bii) was not simply intracellular HGF released during the lysis steps, we sought to separately quantify HGF concentrations in samples containing lysis buffer. Utilizing the DF beads, we quantified the concentration of HGF present in the cell lysate (cells had been rinsed to remove extracellular HGF prior to lysis) and observed significantly less HGF signal in cell lysate when compared to the signals captured from localized cell-surface sampling (Figure S9), indicating that the majority of the HGF captured using our localized cell-surface sampling method is secreted from the cell. Further, in future work, we will develop gentler methods to remove the bead from the cell surface without lysing the cell. In summary, our DF beads can selectively bind to the surface of a cell and capture signal directly secreted from the cell before the secreted factor is sequestered and lost in bulk solution.

## Conclusion

In this work, we present a new bead-based system that enables *in situ* capture of a secreted molecule (HGF) in the presence of a known neutralization factor (free anti-HGF); through targeted cell-tethering of DF beads, we can capture secreted HGF as it is released from the cell, rescuing signal that is lost in traditional supernatant analysis due to HGF binding to free anti-HGF in the bulk solution (Figure 4). MagPlex beads were chosen to enable future development and multiplexing of different analytes and cell types, as they are commercially available and compatible with a wide range of antibodies, allowing easy adaptation of this technology without the need for specialized fabrication or experimental protocols. Further development of this platform will focus on selective bead detachment methods and optimizing antibody ratios to improve binding capabilities and permit continuation of cultures after bead detachment. Future application of this technology includes deploying this method in complex biological system containing multiple cell types (e.g., mixed cocultures of mammalian cells, multikingdom culture containing microbial and mammalian cells, and *ex vivo* tissue slices) to demonstrate our method’s potential to selectively target a specific cell type in a complex environment containing multiple cell types, as well as complex biological samples (e.g., blood) to capture short-lived factors for physiological analysis.

## Supporting Information

The Supporting Information for this work includes supplementary data further characterizing the DF bead functionalities within the model system discussed in this work, as well as control experiments supporting the work presented in Figures 1-4. Additionally, the complete experimental materials are included in the Supporting Information.

## Acknowledgements

This work was supported through an award from the Kavli Microbiome Ideas Challenge, a project led by the American Society for Microbiology in partnership with the American Chemical Society and the American Physical Society and supported by The Kavli Foundation (preliminary experiments); by the National Science Foundation Graduate Research Fellowship Program under Grant No. DGE-1256082 (SBB), and NIH1R35GM128648 (ABT, EB, TLVN). Any opinions, findings, and conclusions or recommendations expressed in this material are those of the author(s) and do not necessarily reflect the views of the National Science Foundation or the National Institutes of Health. We graciously thank Dr. Angela Wilson (University of Washington, Department of Pathology) and Richard Lawler (Fred Hutchinson Cancer Research Center) for all their training and guidance on Luminex assays and instrumentation.

## Conflicts of Interest

The authors acknowledge the following potential conflicts of interest in companies pursuing open microfluidic technologies: EB: Tasso, Inc., Salus Discovery, LLC, and Stacks to the Future, LLC; ABT: Stacks to the Future, LLC. However, these companies are not related to the methods presented in this paper.

## Notes

### Competing Interest Statement

The authors have declared no competing interest.

