## Supplementary material for "Localized cell-surface sampling of a secreted factor using cell-targeting beads": SI

Supporting Information:

**Materials and Methods**

**Supplementary Figures**

Figure S1: CD90 expression
Figure S2: Validation of dual-functionalized beads
Figure S3: Bead-cell binding validation
Figure S4: Phase contrast imaging of bead-cell binding validation
Figure S5: High magnification (20X) phase contrast imaging
Figure S6: Validation of CD90$^+$ cells selectively binding anti-CD90, anti-HGF DF beads
Table S1: Total bead counts for bead-cell binding validation
Figure S7: Specific cell-targeting capabilities
Table S2: Experimental reproducibility of bead-based methods
Figure S8: Transwell control experiment
Figure S9: HGF concentration within cell lysate

**References**

### Materials and Methods

*Reagents and Materials*

Luminex kits (Product # HAGP1MAP-12K) containing wash buffer, assay buffer, secondary detection antibody, mono-functionalized beads, recombinant human hepatocyte growth factor (HGF), and streptavidin-R-phycoerythrin conjugate (SAPE) were purchased from EMD Millipore (Burlington, MA) and used for anti-HGF/HGF Assay experiments (details below). Clear tissue culture treated (TCT) 96-, 24-, and 12-well plates (Corning #3596, #3526, and #3515, respectively) and black TCT 96-well plates (Corning #3603) were purchased from Corning (Corning, NY). Falcon Transwell 24-well plate inserts (Corning #353104) were purchased from Fisher Scientific (Hampton, NH). Antibody coupling kits (Product #40-50016) were purchased from Luminex Corporation (Austin, TX).

*Cell Culture*

Normal Human Dermal Fibroblast neo-natal cells (NHDFn) (ATCC, Manassas, VA) were cultured, passaged, and seeded in fibroblast basal medium supplemented with a low-serum growth kit (ATCC, #PCS-201-030 and #PCS-201-041) (fibroblast media). 24 hours after seeding cells in well plates or Transwell inserts, media was replaced with Endothelial Cell Growth Medium-2 BulletKit (EGM-2) (Lonza, Basel, Switzerland) supplemented with 10% heat-inactivated (HI) fetal bovine serum (FBS) (see below); media was changed daily. NHDFn passage numbers ranged from 6-8 for experiments. Human Umbilical Vein Endothelial cells (HUVEC) (Lonza, Basel, Switzerland, #C2517A) were cultured, passaged, and seeded in EGM-2 media. HUVEC passage numbers ranged from 3-5 for experiments. FBS was heat-inactivated following the procedure provided by Serum Source International. Briefly, FBS was brought to room temperature and placed in a water bath heated to 56 ˚C for 30 minutes. HI-FBS was then cooled to room temperature before being stored at -80 ˚C until use.

*CD90 Expression by Normal Human Dermal Fibroblast neo-natal Cells (NHDFn)*

Expression of CD90 by NHDFn cells was validated through standard immunocytochemistry techniques. NHDFn cells were cultured in a black TCT 96 well plate for 3 days; on Day 3 post seeding, cells were fixed for 10 minutes with 4% paraformaldehyde, permeabilized with 0.2% Triton X-100, and then blocked with 3% bovine serum albumin (BSA). After blocking, primary anti-CD90 monoclonal antibody (clone AF-9, Abcam, #ab23894) (1:50 dilution) was added and incubated overnight at 4°C; goat anti-mouse secondary polyclonal antibody conjugated with Alexa Fluor-488 (1:200 dilution) (Jackson ImmunoResearch Laboratories Inc., #115-545-166) was then allowed to incubate for 1 hour at room temperature. After washing, DAPI (ThermoFisher, #D1306) (1:200 dilution) was incubated for 5 minutes at room temperature before washing and imaging (Figure S1). HUVECs were simultaneously cultured as described above to serve as a negative control. HUVECs were stained with CellTracker Blue (ThermoFisher, #C2110) for 1 hour before seeding according to the manufacturer's protocols. Fixing and staining protocols outlined above were followed before imaging (Figure S1).

*Bead Functionalization*

MagPlex microspheres (region 34) were purchased from Luminex Corp. and functionalized with anti-human hepatocyte growth factor (anti-HGF) monoclonal antibody [(clone 24516) (R&D Systems, Inc., Minneapolis, MN)] and anti-human cluster of differentiation 90 (anti-CD90) monoclonal antibody (clone

AF-9) (Abcam, Cambridge, MA) according to the manufacturer's instructions (Luminex antibody coupling kit, #40-50016).[1] Briefly, carboxylated microspheres were first activated with 1-ethyl-3-[3-dimethylaminopropyl] carbodiimide hydrochloride (EDC) and sulfo-N-hydroxysulfosuccinimide (Sulfo-NHS). Once activated, anti-CD90 and anti-HGF antibodies were added at a 1:1 concentration-based ratio to covalently bind to the surface of the activated microspheres. Once coupled, microspheres were rinsed and stored in phosphate buffered saline (PBS) containing 1% bovine serum albumin (BSA) at 4°C until use. Care was taken to protect the microspheres from light to prevent photobleaching during coupling. Microspheres were used as per the manufacturer's instructions and within 6 months of coupling. To validate successful coupling of the antibodies with the microspheres, we performed a secondary antibody labeling and subsequent analysis.[1] Briefly, phycoerythrin-conjugated rat anti-mouse IgG (clone M1-14D12) (ThermoFisher Scientific) was added to the microspheres at a range of concentrations (0.0625 µg/mL to 4 µg/mL), incubated, and analyzed to generate a calibration curve as per the manufacturer's instructions (Figure S2). Antibodies used for bead functionalization (anti-HGF and anti-CD90) were validated by manufacturer's using XYZ method(s).

*Bead-Cell Binding Validation*

To validate the bead-cell binding,_NHDFn cells were passaged, incubated with CellTracker Green for 1 hour at room temperature, washed with 1X PBS, and seeded in a black TCT 96 well plate at a density of $2.6x10^4$ cells/mL and cultured for 3 days; cells were seeded in fibroblast media and replaced with EGM-2 media for the remainder of the experiment. As a negative control, HUVECs were passaged, incubated with CellTracker Blue for 1 hour at room temperature, washed with 1X PBS, and seeded at a density of $2.6x10^4$ cells/mL; cells were cultured for 3 days in EGM-2 media. Dual-functionalized (DF) beads (coupled with both anti-CD90 and anti-HGF antibodies) at a concentration of 50 beads/µL were added to the cultures on Day 3, post-seeding. An additional negative control was used in which different dual-functionalized beads [(coupled with both anti-CD64 (cluster of differentiation 64) and anti-MMP12 (matrix metallopeptidase 12) antibodies) (Abcam, #ab119843 and #ab52897, respectively)] were added to separate wells containing NHDFn cultured in the same manner as stated above. Plates were shaken at 100 rpm for 1 minute then placed on a plate magnet (Stemcell Technologies, #18102) for 1 minute; this was repeated twice more for a total of three rounds followed by a 2-hour incubation. After incubation, wells were washed with EGM-2 media 3 times. Wells were imaged before washing wells and after washing wells (Figure S3-S6). To demonstrate cell-targeting ability, a device (Monorail2) was used to pattern NHDFn and HUVECs in separate regions of the well plate.[2] Full device operation and protocol is provided in Day et al.[2] Briefly, a Monorail2 device was placed into a 24-well plate and a 1.5 wt% low gelling temperature agarose pre-gel solution (Sigma-Aldrich, #39346-81-1) was flowed to create a hydrogel wall; gel was cooled at room temperature until solidified. 1X PBS was loaded into wells for 24 hours before use in experiments. Prior to seeding, NHDFn were incubated with CellTracker green dye for 1 hour at room temperature and HUVECs were incubated with CellTracker blue dye for 1 hour at room temperature (following dilution instructions provided) and washed with 1X PBS. Both cell types were seeded at a density of $2.6x10^4$ cells/mL and cultured for 3 days in EGM-2 media. On Day 3 post-seeding, Monorail2 devices were removed and dual-functionalized beads (coupled with both anti-CD90 and anti-HGF antibodies) at a concentration of 50 beads/µL were added to the cultures. Experimental workflow outlined above (shaking, washing, incubation, and imaging steps) was followed (Figure S7).

*Bead-Analyte Binding Validation*

A model capture sandwich immunoassay was performed using both dual-functionalized beads (anti-CD90 and anti-HGF antibodies) and commercial mono-functionalized kit beads (Product # HAGP1MAP-12K) (anti-HGF antibody only) according to manufacturer's instruction. Briefly, diluted beads (50 beads/µL) were incubated with recombinant human HGF protein [(clone #24516) (R&D Systems, Minneapolis, MN)] at concentrations ranging from 0 to 20,000 pg/mL (standard curve range according to each commercial kit). After incubation, beads were washed and detection antibody (biotinylated anti-human HGF polyclonal antibody, #BAF294, R&D Systems) was added to label the captured HGF. After binding of the detection antibody, streptavidin-R-phycoerythrin conjugate (SAPE) (ThermoFisher, #S866) was added and the corresponding fluorescence of the beads was analyzed. A five-parameter logistic fit (5PL) was used to fit calibration curves based on kit protocols. Calibration curves from each experiment, for both mono- and dual-functionalized beads, are shown in Figure 2. Curves were used to determine HGF concentration for each independent experiment. Limit of detection (LOD) was calculated using the standard definition of three times the background signal for each curve. Per manufacturer's instructions, if a well did not contain sufficient bead count, it was not analyzed and was excluded from further data analysis (Figure 2C Experiment 3 20,000 pg/mL HGF standard, Figure 4Bii Experiment 3 control condition, Figure S9 Experiment 3 control condition).

*Cell-secreted HGF Assay*

Media preparation: Two different media were used, one control and one treated. The control media consisted of EGM-2 supplemented with 10% HI-FBS and 1X PBS in place of antibody (control media). The treated media consisted of EGM-2 supplemented with 10% HI-FBS and free anti-HGF [(clone 24516) (R&D Systems, Inc., Minneapolis, MN)] in 1X PBS for a final concentration of 10 ng/mL free anti-HGF (treated media). Media were made fresh on the day of the experiment.

Localized cell-surface sampling: NHDFn cells were cultured in a clear TCT 96 well plate as described above. On Day 3 post seeding, cells were washed once with media; 75 µL control (- free anti-HGF) or treated (+ free anti-HGF) media and 50 µL of diluted dual-functionalized (anti-HGF and anti-CD90) beads (100 beads/µL) in control media were added and incubated with the cells for 2 hours. To distribute added beads within a well, the plate was shaken at 100 rpm for 1 minute, then placed on a plate magnet (Stemcell Technologies, #18102) for 1 minute; this was done for a total of 3 rounds in the beginning of the incubation period. After 2 hours, well contents were washed twice with 125 µL control media to remove any beads not bound to the cell surfaces; 50 µL of lysis buffer (10 mM Tris-HCl, 1 mM EDTA, 1% Triton X-100, 0.1% SDS, 140 mM NaCl) was then added and incubated for 15 minutes to detach beads from the cell surface. A mini cell scraper (ABI Scientific Inc., catalog #MCS-200) was used 5 minutes into the 15-minute lysis step to break up cellular debris during the lysing period. The plate was then placed on a magnet, and beads were washed with control media 3 times before being resuspended in 50 µL control media, collected, and added to assay plate; this resuspension was used as the final sample with no additional beads added and utilized for the Luminex assay. The assay was continued using the Luminex kit and procedure.[1] Briefly, samples were incubated for 16-20 hours on an orbital shaker at 4˚C. Beads are washed with wash buffer and secondary detection antibodies are added for a 1-hour room temperature incubation. SAPE is then added for an additional 30 minutes. Additional wash steps remove any unbound antibodies and SAPE before beads are resuspended in 1% BSA for analysis with FLEXMAP 3D System instrument.

Traditional bulk supernatant sampling: Cell culture methods were the same as described in the above paragraph. On Day 3 post seeding, cells were washed once with media; 75 µL control (- free anti-HGF) or treated (+ free anti-HGF) media and 50 µL of control media were added and incubated with the cells for 2 hours. Cell culture supernatant samples were collected from each well and transferred to new wells in a 96 well plate to be analyzed with mono-functionalized beads. The assay was continued using the Luminex kit and procedure.[1] Briefly, mono-functionalized (anti-HGF only) beads are added to supernatant samples to incubate for 16-20 hours on an orbital shaker at 4˚C for HGF analyte capture. [Note: while dual-functionalized beads (anti-HGF and anti-CD90) described in the prior paragraph undergo the same process, it is important to note that analyte capture is completed prior to addition onto the assay plate for the dual-functionalized beads.] Beads are washed with wash buffer and secondary detection antibodies are added for a 1-hour room temperature incubation. SAPE is then added for an additional 30 minutes. Additional wash steps remove any unbound antibodies and SAPE before beads are resuspended in 1% BSA for analysis with FLEXMAP 3D System instrument.

*Transwell assays:* Inverted Transwell inserts were placed into a clear TCT 12 well plate after which a 50 µL fibroblast cell suspension (2.6x$10^4$ cells/mL) was placed on top of the Transwell inverted insert (in the pore membrane area); cells were incubated for 2 hours to allow cell adherence before Transwell inserts were flipped and placed correctly into a clear TCT 24 well plate with 500 µL control media in the basal side (Figure S8 A). Additionally, in other wells, cells were also cultured on the bottom of a clear TCT 24 well plate at the same density (Figure S8 B). Cells were cultured as described above for the remainder of the experiment. On Day 3 post seeding, 50 µL diluted beads (100 beads/µL) and 50 µL +/- free anti-HGF media were added to the apical side of the Transwell inserts while 500 µL +/- free anti-HGF media was added to the basal side for a 2-hour incubation. Beads were collected from the apical side of the Transwell inserts and washed twice before being resuspended in 100 µL control media and added to assay plate; this resuspension was used as sample with no additional beads added. The assay was continued as previously described above using the Luminex kit and procedure. The data are presented in Figure S6.

*Instrumentation*

Analysis for all bead-based assays (MF and DF beads) were performed on a FLEXMAP 3D System (Luminex Corp.) in the Immune Monitoring Lab (Fred Hutchinson Cancer Research Center, Seattle, WA). The microspheres were analyzed with the following settings: 50 beads, 50 events/bead, 75 µL, bead region 45 (commercial kit mono-functionalized beads) or 34 (dual-functionalized beads), 5000 – 30000 gate, 60 second time out. Fluorescent images of the cells and beads were obtained using a Zeiss Axiovert 200 and an Axiocam 503 mono camera (Carl Zeiss AG, Oberkochen, Germany). Data and statistical analyses were completed using Prism (Graphpad, San Diego, CA) software. All cell and bead images were processed using FIJI image software (ImageJ, NIH).

**Supplementary Figures:**

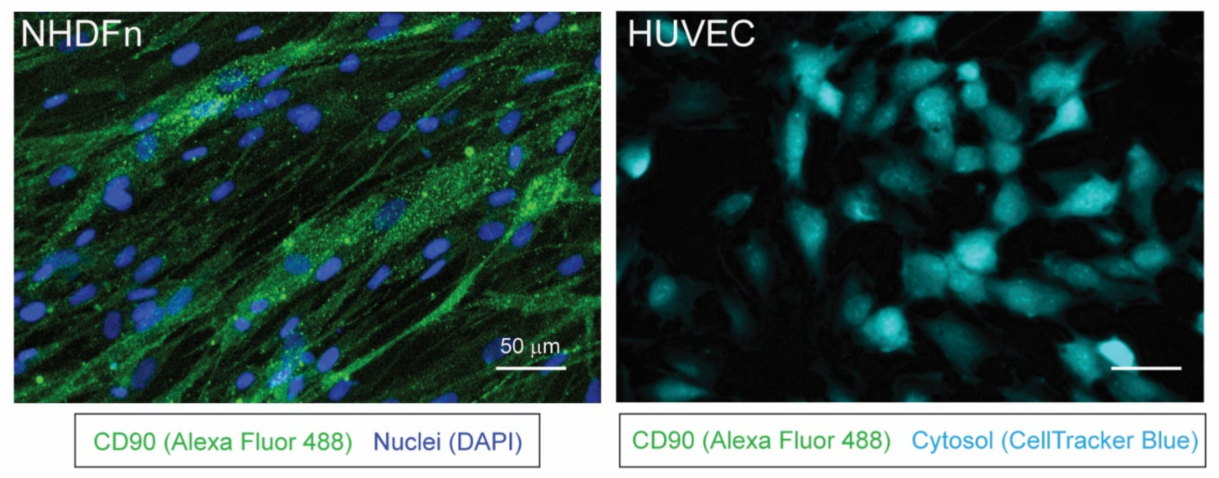


**Figure S1**: Validation of CD90 expression by NHDFn cells using immunocytochemistry. CD90 expression (green) was confirmed in NHDFn ($CD90^+$ cells) using standard immunocytochemistry techniques. HUVECs ($CD90^-$ cells) were stained with CellTracker Blue (light blue) and did not show CD90 expression (no green signal observed). Scale bar is 50 µm.

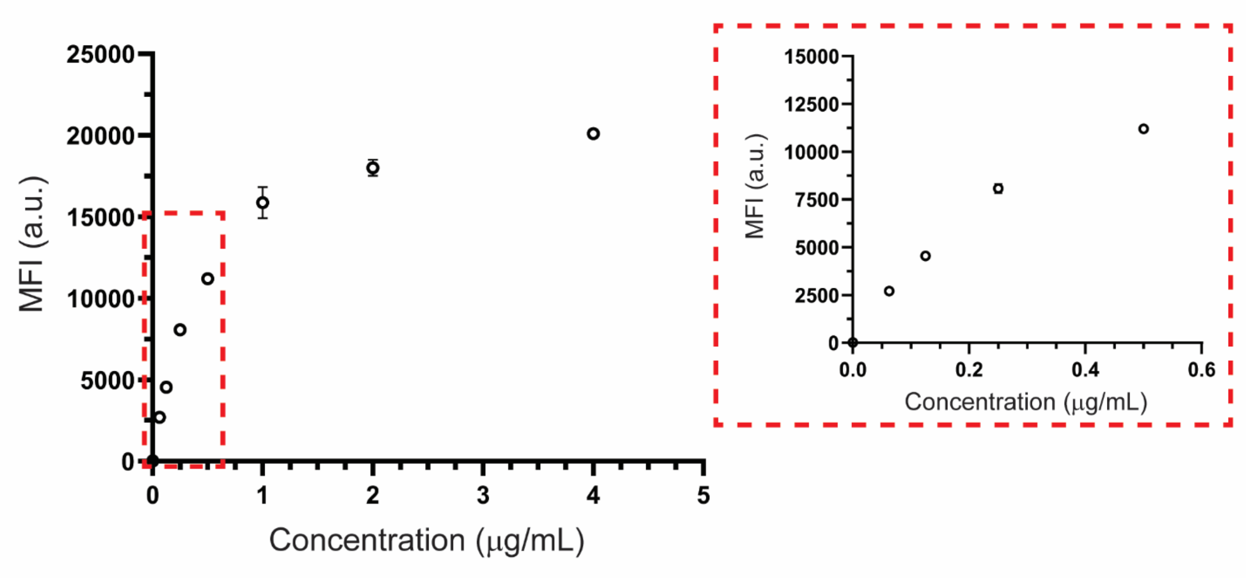


**Figure S2**: Validation of antibody functionalization on MagPlex bead surface. Antibodies (CD90 and HGF) were covalently coupled to the bead surface and then incubated with a range of fluorescent secondary antibody concentrations to demonstrate successful coupling of antibodies to bead surface (see Materials and Methods for details). The plot reflects an expected response based on the manufacturer's (Luminex) protocol.[1] Data points are the average of duplicate measurements with error bars representing standard deviation. The graph on the right is an inset of the 0-0.5 µg/mL concentration range.

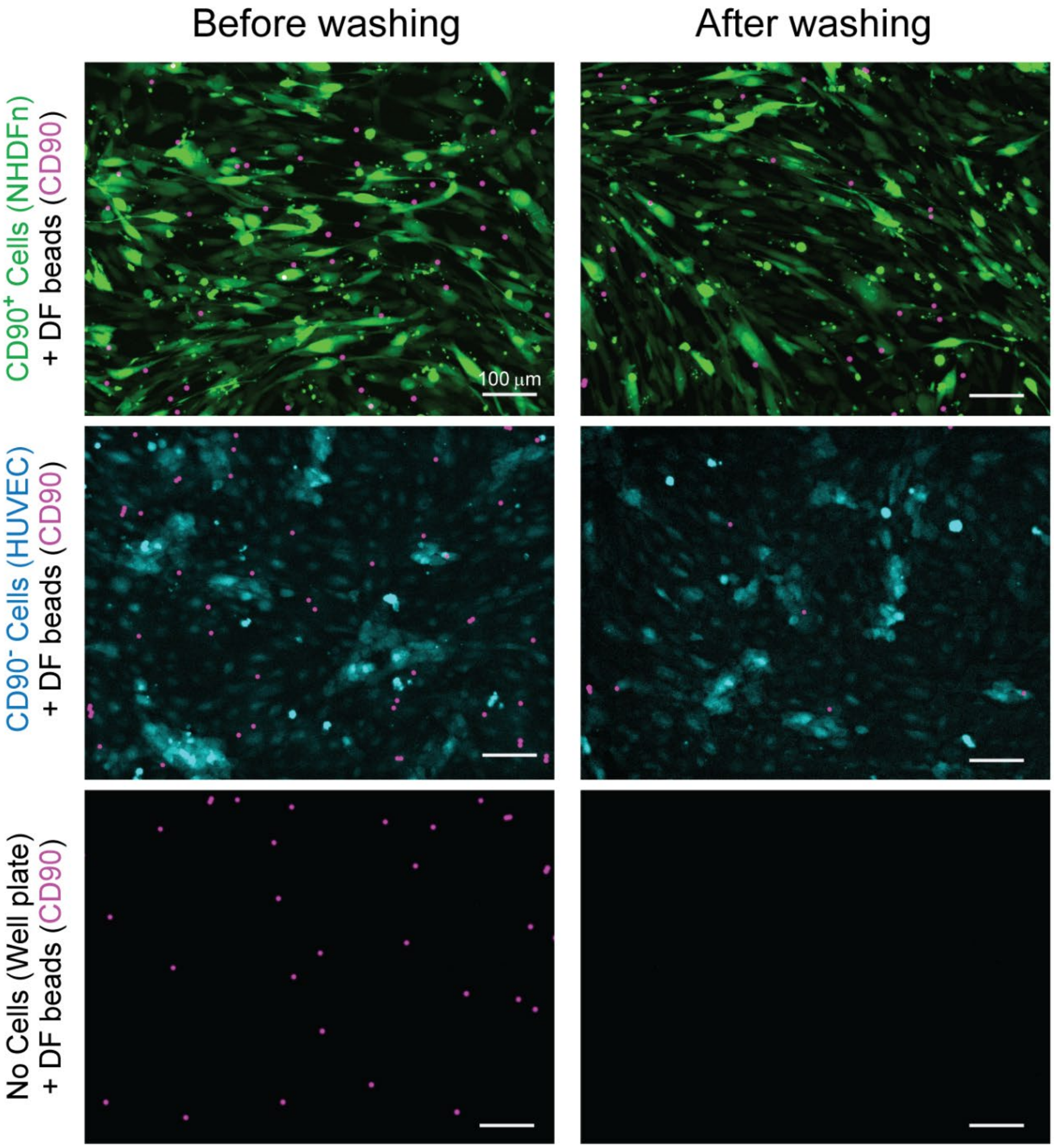


**Figure S3**: Validation of dual-functionalized (DF) bead binding to the surface of targeted $CD90^+$ cells (NHDFn). Representative images from n=3 independent experiments demonstrating DF bead [(anti-CD90, anti-HGF) (pink)] tethering to NHDFn cells (green) after multiple washing steps. Bead retention was also quantified in two negative control conditions where bead retention after washing is not expected: anti-CD90, anti-HGF DF beads with $CD90^-$ cells [(HUVECs) (blue)]; anti-CD90, anti-HGF DF beads in a well plate with no cells. All negative conditions showed negligible bead retention on the cell/well plate surface. Images are representative of 3 independent experiments. Bead retention data is quantified in Figure 3C and Table S1. Scale bar is 100 µm.

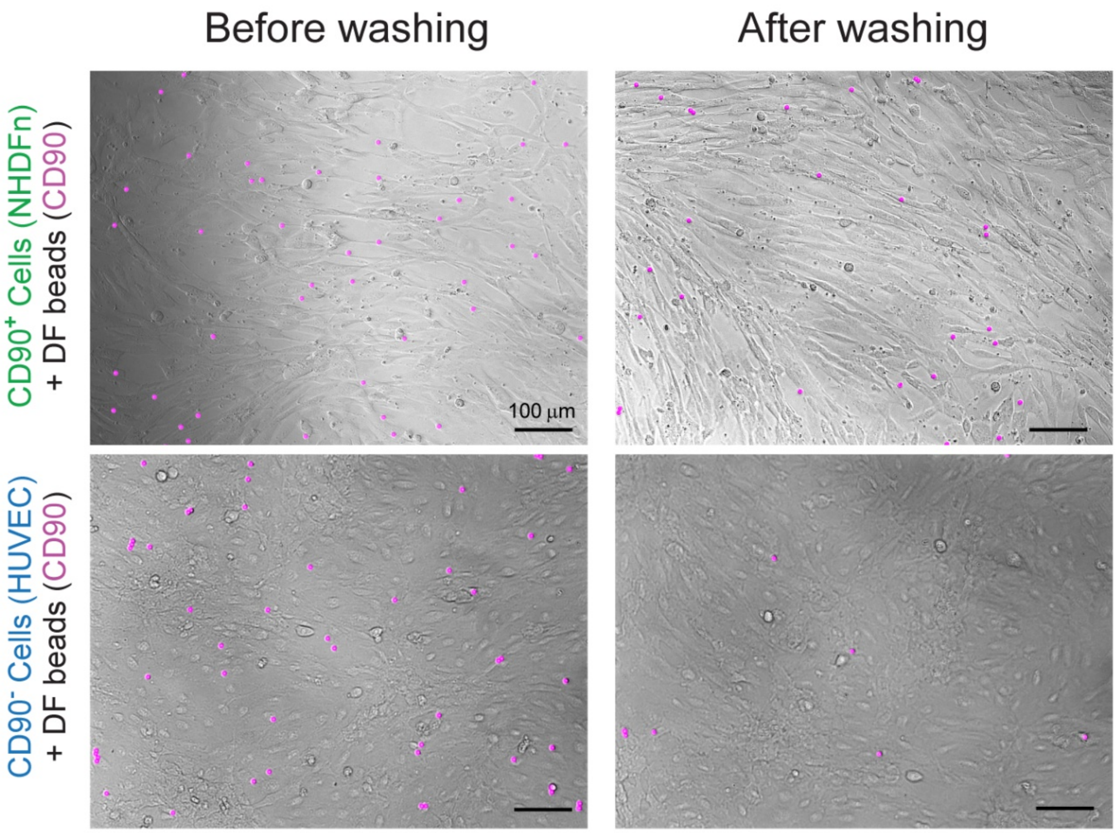


**Figure S4**: Phase contrast imaging of dual-functionalized beads binding to the surface of targeted CD90$^+$ cells (NHDFn). Here we can see DF beads remaining after washing are bound to the surface of targeted CD90$^+$ cells. In the non-targeted CD90$^-$ cell condition, remaining beads are mostly on the well plate surface. Fields of view correspond to the images shown in Figure S3 and were taken at 10X magnification. A fluorescence microscopy overlay of the DF beads (pink) was used for better visualization. Higher magnification (20X) images are shown in Figure S5. Scale bar is 100 µm.

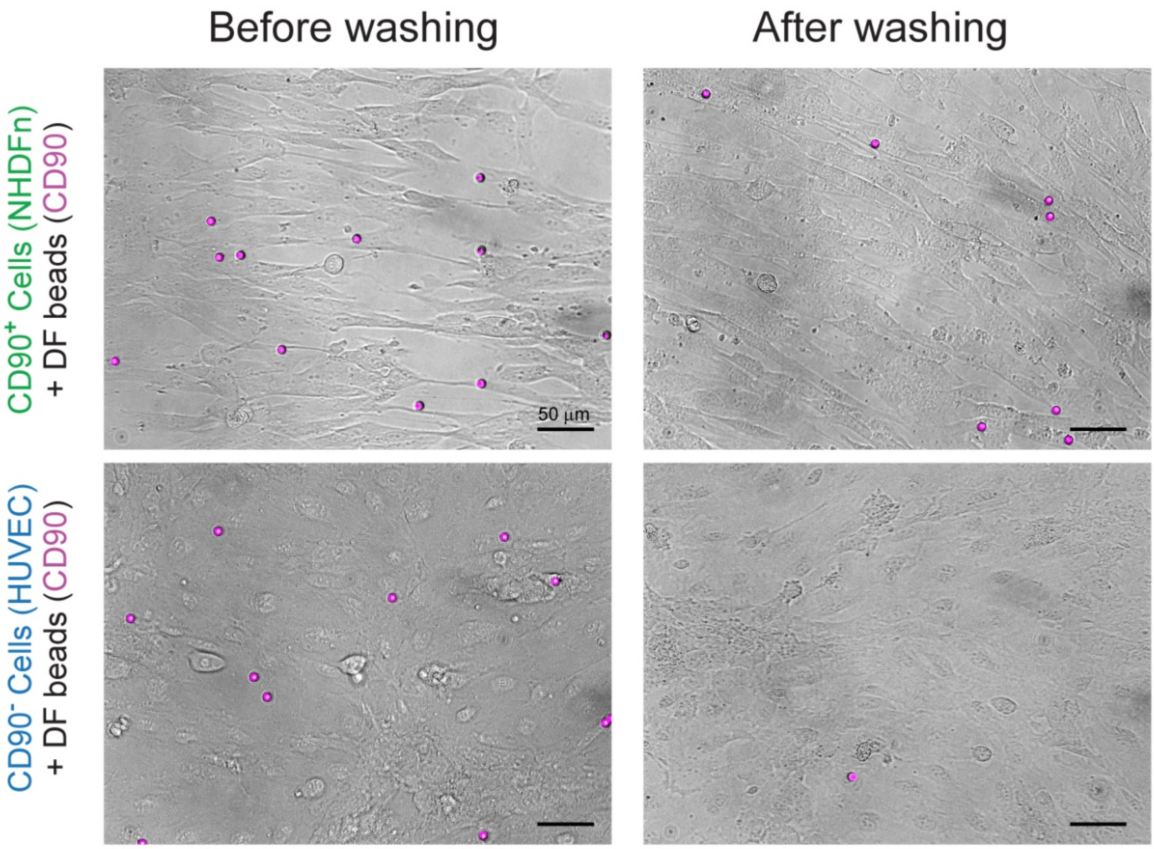


**Figure S5**: 20X magnification phase contrast imaging of dual-functionalized beads binding to the surface of targeted $CD90^{+}$ cells (NHDFn). Fields of view are insets of the images shown in Figure S4. A fluorescence microscopy overlay of the DF beads (pink) was used for better visualization. Scale bar is 50 µm.

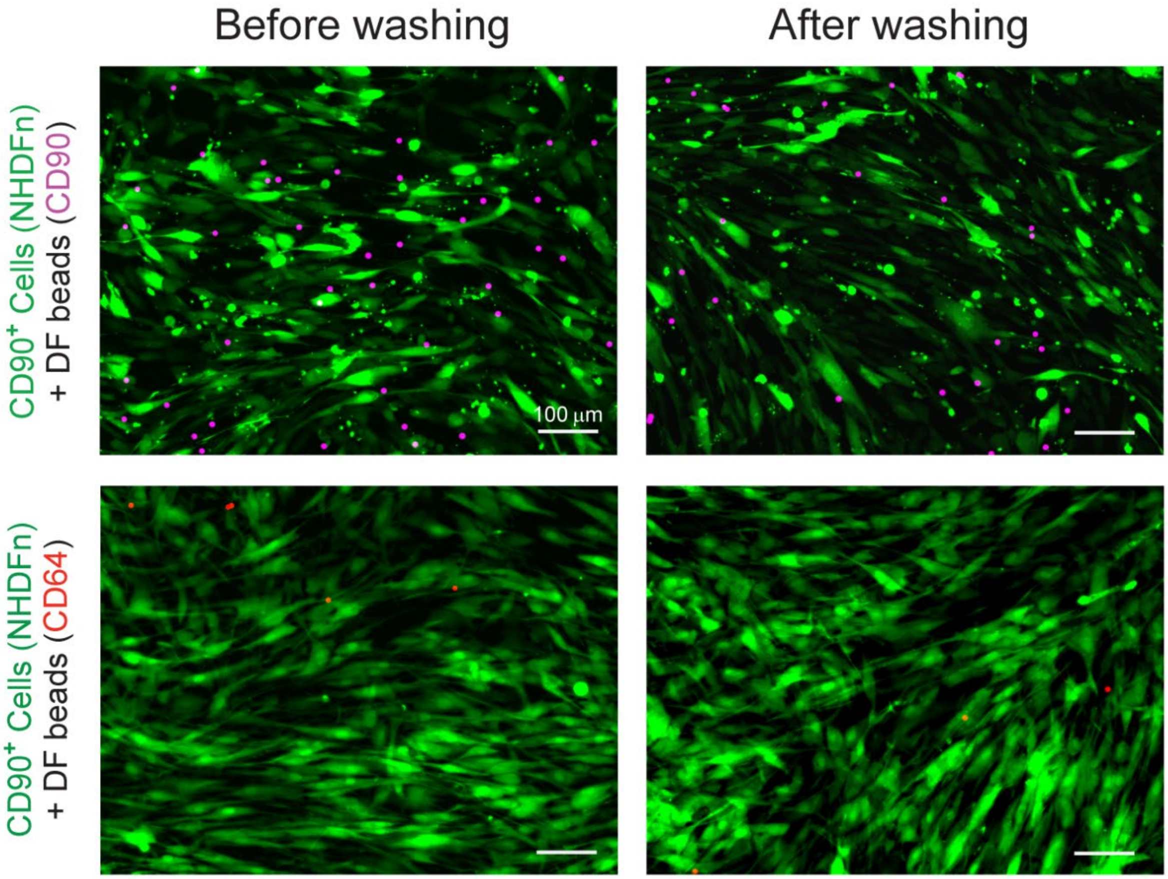


**Figure S6**: Validation of CD90$^+$ cells (NHDFn) selectively binding anti-CD90, anti-HGF DF beads. Representative images from n=3 independent experiments demonstrating DF bead [(anti-CD90, anti-HGF) (pink)] tethering to NHDFn cells (green) after multiple washing steps. Bead retention was also quantified in a DF bead negative control condition using beads functionalized with anti-CD64 and anti-MMP12 (red). Bead retention on NHDFn of anti-CD64, anti-MMP12 after washing is not expected. Images and bead quantification data showed negligible bead retention on the cell/well plate surface (Table S1). Scale bar is 100 µm.

| Condition | Before washing (average ± SD) | After washing (average ± SD) |
|---|---|---|
| $CD90^+$ Cells (NHDFn) + DF beads (CD90) | 29.4 ± 6.6 | 21.0 ± 4.7 |
| $CD90^+$ Cells (NHDFn) + DF beads (CD64) | 8.4 ± 2.8 | 1.6 ± 1.3 |
| $CD90^-$ Cells (HUVEC) + DF beads (CD90) | 29.3 ± 3.6 | 3.3 ± 2.4 |
| No Cells (Well plate) + DF beads (CD90) | 27.6 ± 6.2 | 1.2 ± 1.5 |

**Table S1**: Total bead counts for bead-cell binding validation. The table shows the average bead count ± standard deviation (SD) for the representative images used in Figure 3, S3, and S6. Each condition included n=9 images for n=3 independent experiments.

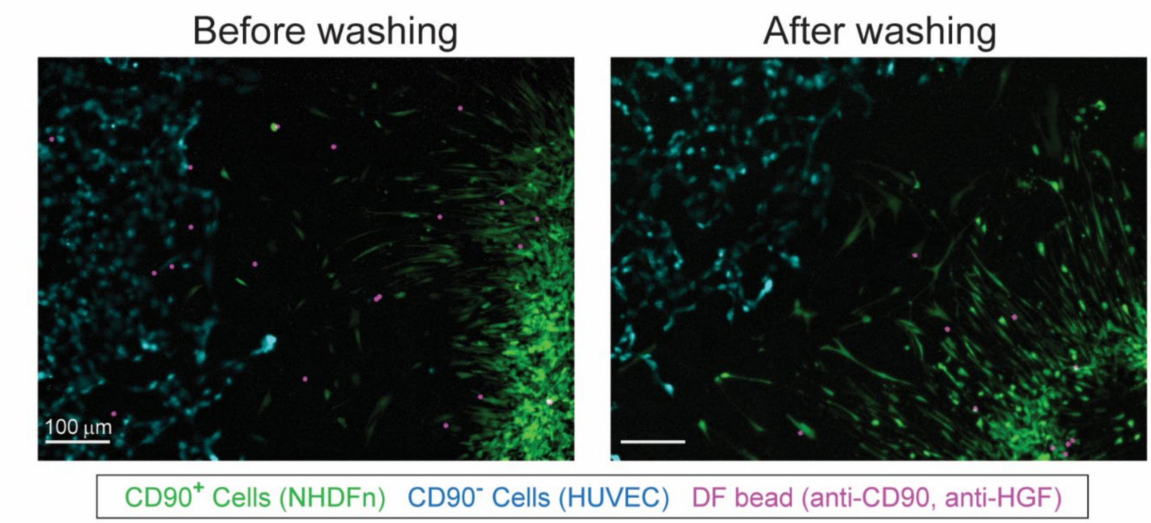


**Figure S7**: Dual-functionalized (DF) beads (anti-CD90, anti-HGF) remain tethered to targeted $CD90^+$ cells (NHDFn) in coculture with $CD90^-$ cells (HUVEC). DF beads (pink) were added to a patterned coculture of NHDFn (green) and HUVEC (blue) cells for a 2-hour incubation followed by multiple wash steps. Beads remain tethered to $CD90^+$ cells (green), while being removed from $CD90^-$ cells (blue) after washing. Scale bar is 100 μm.

| | Control (- anti-HGF) | | Treated (+ anti-HGF) | |
|---|---|---|---|---|
| | MF bead | DF bead | MF bead | DF bead |
| Experiment | RSD (%) | | | |
| 1 | 37.6 | 14.4 | 28.9 | 11.6 |
| 2 | 14.3 | 8.6 | 15.7 | 12.5 |
| 3 | 7.7 | 1.1 | 12.6 | 11.7 |
| 4 | 61.4 | 4.6 | 53.9 | 13.2 |

**Table S2**: Experimental reproducibility of mono- and dual-functionalized (MF and DF, respectively) bead methods. The table includes the relative standard deviation (RSD) for each experiment plotted in Figure 4. More variability exists within the MF bead system (traditional method), while the DF bead system (our new method) is consistently below 15%.

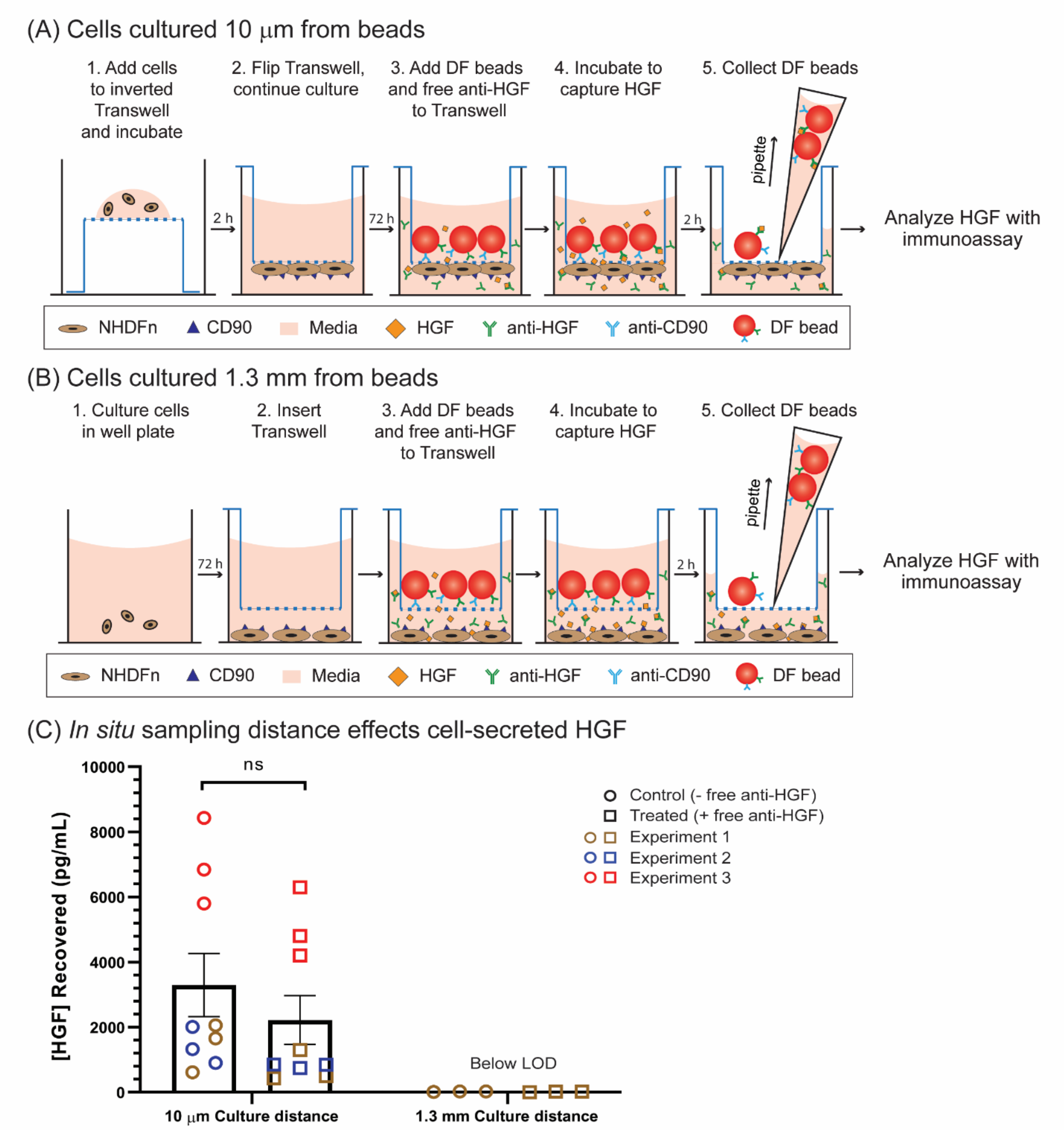


**Figure S8**: Increased *in situ* HGF recovered concentration results from bead proximity to cells. Schematic showing steps for culturing cells (A) 10 μm from beads and (B) 1.3 mm from beads. In both conditions, DF beads are spatially separated from cells and are not attached to the cells. These two distances were chosen to probe the effect of bead sampling distance on HGF capture. When beads were cultured 10 μm from cells there was (C) no statistical significance between the control (- free anti-HGF) and treated (+ free anti-HGF) groups. This same trend was observed when beads were tethered to the cell surface via CD90 (see Figure 3 in the manuscript). In contrast, beads cultured 1.3 mm from the cells resulted in signals below the assay limit of detection (LOD) in both conditions. Three independent experiments with three replicates were performed for each experiment. Error bars are SEM. Unpaired, parametric t-test.

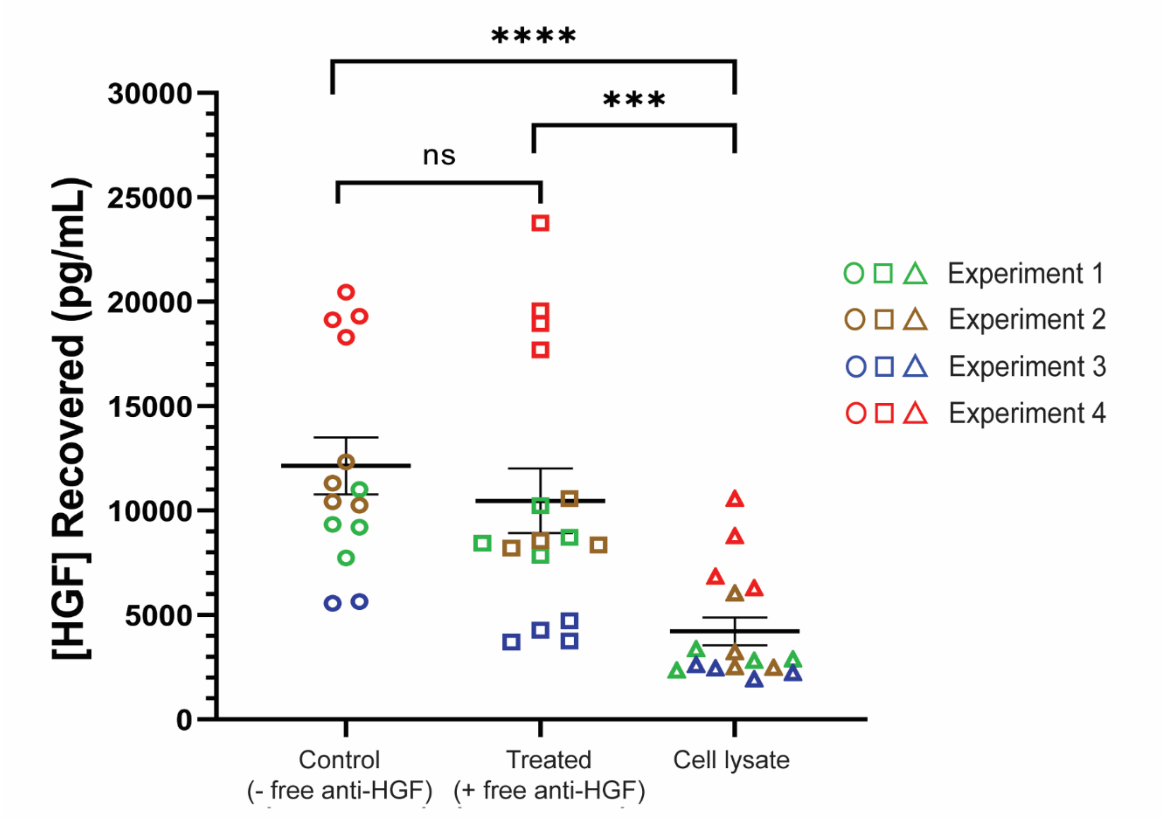


**Figure S9**: Quantification of HGF in cell lysate from the cell lysis step to remove DF beads tethered on cell surface. To determine the contribution of intracellular HGF to total recovered HGF signal, we quantified the amount of HGF present during the 15-minute lysing period used in the localized cell-surface sampling method workflow by simultaneously adding DF beads and lysis buffer to cells. While signal is observed in the cell lysate, the HGF concentration is significantly less than the recovered HGF from either localized cell-surface sampling condition (control and treated). Control and treated conditions were replotted from Figure 3. Each data point plotted represents one technical replicate, with typically 4 replicates per independent experiment. Error bars are SEM. Unpaired, parametric t-test. *** $p < 0.001$, **** $p < 0.0001$.
